# From Michaelis–Menten parameters to microscopic rate constants: an inversion approach for enzyme kinetics

**DOI:** 10.1101/2025.10.23.684102

**Authors:** Silvia Berra, Sara Sommariva, Michele Piana, Giacomo Caviglia

## Abstract

A basic model of enzyme kinetics is the enzyme-catalyzed reaction, in which a single enzyme *E* binds a substrate *S* to form a complex *C*, which subsequently releases a product *P* and regenerates the enzyme. According to the mass-action law, this process can be described by a system of four ordinary differential equations for the four unknown concentrations, governed by the microscopic rate constants: forward (*k*_*f*_), reverse (*k*_*r*_), and catalytic (*k*_cat_). Under suitable assumptions, the full system reduces to the classical Michaelis-Menten model, depending on two experimentally measurable parameters: the half saturation constant (*K*_*m*_), and the limiting rate *V*_max_. This paper addresses the inverse problem of recovering *k*_*f*_ and *k*_*r*_ from *K*_*m*_ and *V*_max_ (or equivalently *k*_cat_, retrievable from *V*_max_). The proposed method exploits a new identity satisfied by any substrate concentration *s*(*t*) in the full mass-action system, which provides the starting point for the inversion procedure. Here, a detailed presentation of an algorithm for reaching this goal is provided, along with an assessment of reconstruction accuracy and an outline of simulations and applications. Beyond enzyme kinetics, the approach supports the construction of novel chemical reaction networks, with potential applications to modeling complex biochemical pathways involved in diseases such as cancer.

## Introduction

It is well known that enzymes play a fundamental role in systems biology, being essential in processes such as cell metabolism, signal transduction, and regulation [1, 2, 3]. Similarly, malfunctions of enzymatic systems may be associated with diseases, including certain types of cancer [4]. Enzymatic activity consists of catalyzing reactions by binding substrates and promoting their conversion into products [5]. A simple model of such a reaction involves a single enzyme (*E*) that binds a substrate (*S*) to form a complex (*C*), which subsequently releases a product (*P*) while regenerating the enzyme (*E*) [6, 5]. This process is represented schematically by the reactions

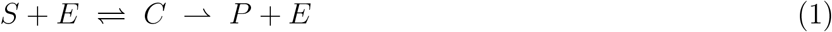

According to the mass-action law, the mathematical description of this system consists of four ordinary differential equations (ODEs) for the concentrations of the involved chemical species. Such equations depend on three microscopic parameters: the forward (association) rate constant *k*_*f*_, the reverse (dissociation) rate constant *k*_*r*_, and the catalytic rate constant *k*_cat_ [5]. Determining the time evolution of the concentrations from the given ODE system, once fixed initial conditions and rate constants, represents a standard example of direct problem.

Under the quasi-stationarity assumption for *C*, the enzyme-catalyzed reaction model (1) can be approximated by the Michaelis–Menten (MM) kinetics [2, 5, 7, 8]. In this framework, the dynamics reduce to a single nonlinear ODE for the substrate concentration *s*, depending on the two MM parameters: the Michaelis constant *K*_*m*_, and the limiting rate *V*_max_. If the total enzyme concentration *e*_0_ is known, the related catalytic constant is given by *k*_cat_ = *V*_max_*/e*_0_ [8]. As for the product *P*, its concentration is directly related to that of the substrate [9, 5, 10]. Beyond its formal simplicity and reasonable accuracy, the MM formulation has the great advantage that a wide variety of experimental and analytical methods have been developed to estimate its parameters under different experimental designs and data-analysis approaches (see, e.g., [9, 2, 8, 3]). This makes MM formulation a cornerstone in quantitative enzyme kinetics [11, 12].

Classical deductions of the standard MM equation, achieved by simplifying the mass-action system, provide the explicit expression for MM parameters in terms of the microscopic rate constants [5, 10, 13]. In contrast, it seems that only one general method for reconstructing *k*_*f*_ and *k*_*r*_ from the MM parameters is available - the so-called Lambda approximation method [14], which however shows high arbitrariness and a significant reconstruction error. Yet, knowledge of the rate constants, when available, provides valuable insight into the dynamics of enzymatic reactions, e.g. by revealing the association and dissociation properties of the substrate–enzyme complex. Their availability also enables the direct application of the mass-action law, leading to an accurate description of enzyme kinetics. From a formal viewpoint, the mass-action formulation is particularly convenient when the reaction is embedded in a larger biochemical network composed of further mass-action reactions. In this setting, the resulting ODE system has a polynomial structure, which facilitates the study of equilibria [15, 16]. More generally, constructing chemical reaction networks based on either experimentally derived or computationally deduced rate constants is essential for simulating the behavior of complex cellular pathways in different scenarios: physiological, altered, or drug-treated. Such networks find applications in diverse biomedical contexts, such as oncology, where they have been used to study cancer-related pathways [17, 18, 19]These considerations highlight the importance of having a simple procedure to estimate rate constants from experimentally accessible data, to both improve the quantitative characterization of enzymatic processes and support more accurate modeling strategies in systems biology.

The present work addresses the determination of the microscopic rate constants *k*_*f*_ and *k*_*r*_ under the assumption that the conditions for the MM approximation are satisfied, and that MM parameters are available. In this setting, the standard MM ODE, written in terms of *K*_*m*_ and *k*_cat_, admits a solution *s*^*a*^ which can be employed as estimate of the substrate concentration *s*. Starting from *s*^*a*^, the unknown rate constants *k*_*f*_ and *k*_*r*_ of the original ODE system are reconstructed: from a mathematical perspective, this constitutes an inverse problem [20], where *s*^*a*^ plays the role of the experimental known input, and the reconstruction of the original parameters is achieved through an inversion procedure.

The key step of the inversion procedure involves the ODE system associated with the model (1), from which an identity is extracted, which links the rate constants and the approximation *s*^*a*^ of the time-dependent substrate concentration profile *s*. The analysis further incorporates the relation between the rate constants and the Michaelis constant *K*_*m*_, which serves as an a priori constraint and is therefore referred to as the constraint equation. By combining the fundamental and constraint equations, a cost function is obtained, whose minimization yields an estimate for *k*_*f*_ and *k*_*r*_. The inversion procedure is illustrated through a series of simulations. The MATLAB^®^ codes implementing the proposed method are openly available at the GitHub repository https://github.com/theMIDAgroup/CRN_MM2M.

The paper is organized as follows. First, the mathematical model of a simple enzyme-catalyzed reaction is briefly revisited. Next, the new identity relating the microscopic rates to the substrate concentration *s* is deduced, the derivation of the MM equation from the mathematical model is reviewed, and the inverse problem for the rate constants *k*_*f*_ and *k*_*r*_ is formulated. A detailed description of the inversion procedure is provided, along with a presentation of the numerical results and applications. Also the accuracy of the reconstructed microscopic rates is discussed, considering the challenges highlighted by sensitivity analysis. Finally, the results are discussed and our conclusions are offered.

## Material and methods

In this Section, the construction of the standard MM equation from the simple model of enzyme-catalyzed reaction is reviewed, essentially following [5, 10, 13], and a new identity for the substrate concentration *s* is deduced. In doing so, the basic quantities, parameters, and results needed for the formulation of the inverse problem of determining the rates *k*_*f*_, *k*_*r*_ are introduced.

### Simple model of enzyme kinetics

Consider the interaction between an enzyme *E* and a substrate *S*, with *C* denoting the enzyme-substrate complex *SE*, and *P* the reaction product. Their concentrations (nM) are indicated by the lowercase letters *e, s, c, p*, respectively. Assuming that a rapid equilibrium assumption holds [8] and that the product *P* does not bind to the free enzyme *E*, the correspondent reactions can be written as

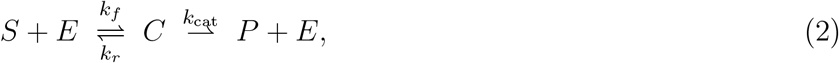

where *k*_*f*_ (nM^−1^ s^−1^) is the forward (association) rate constant, *k*_*r*_ (s^−1^) is the reverse (dissociation) rate constant, and *k*_cat_ (s^−1^) is the catalytic rate constant. The associated system of ODEs for the unknown concentrations is given by

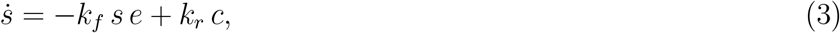

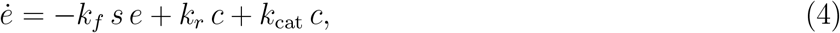

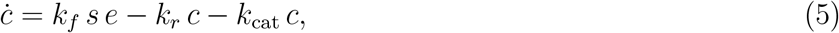

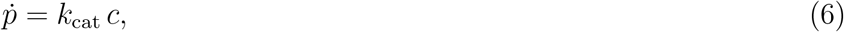

where dependence on time *t* (in seconds s), is not shown explicitly, unless required for clarity, and a superposed dot denotes the derivative with respect to *t*. Equations (3)-(5) form a system of three coupled nonlinear ODEs for the unknowns *s, e, c*. Eq. (6) determines *p* through the integration of *c*. It is reasonable to assume that *c*(0) = *p*(0) = 0, reflecting the fact that the concentrations of *C* and *P* vanish at the initial time. Thus, the only relevant initial conditions are *s*(0) = *s*_0_ and *e*(0) = *e*_0_.

By summing (3), (5), and (6), the conservation law *s* + *c* + *p* = *s*(0) + *c*(0) + *p*(0) = *s*_0_ is obtained. Similarly, summing (4) and (5) yields the further conservation *e* + *c* = *e*(0) + *c*(0) = *e*_0_: the total substrate concentration and enzyme concentration remain constant, i.e., they are not consumed in the reactions.

The conservation law for *e*, written as *e* = *e*_0_ − *c*, can be used to eliminate Eq. (4), thereby reducing the system (3)-(5) to the following set of two ODEs:

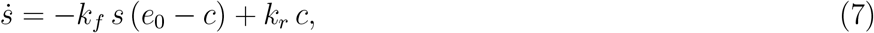

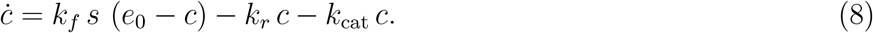

Equations (7) and (8) form a coupled system for the unknowns *s* and *c*. Once *s* and *c* are determined, the concentrations *e* and *p* can be readily obtained. In this sense, the reduced system (7) - (8) and the full system (3)-(6) are essentially equivalent.

### An identity satisfied by substrate concentration

The section is devoted to the introduction of an identity, necessarily satisfied by the solutions of the system (7) - (8), involving only the three rate constants, and the substrate concentration *s*(*t*). This relation plays a crucial role in the formulation and solution of the inverse problem. A representation of *c* in terms of *s* is also found.

The sum of (7) and (8) yields

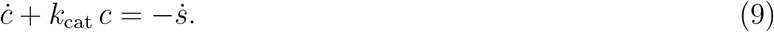

Equation (9) is regarded as a first order ODE for *c*, given 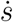. The solution satisfying *c*(0) = 0 is

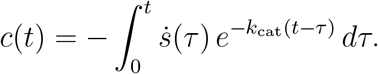

Integration by parts provides an equivalent expression in terms of *s*:

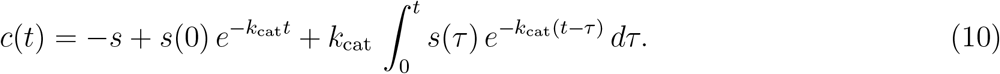

Equation (10) implies that *c* depends on *s*, so that integrating (7) with respect to *t* over [*t*_0_, *t*], with 0 ≤ *t*_0_ *< t*, leads to

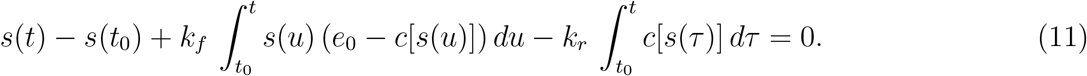

Note that this relation is linear in *k*_*f*_ and *k*_*r*_, since *c* does not depend explicitly on these rates. In the formulation of the inverse problem, the substrate concentration *s* in (10) and (11) is approximated by the solution of the MM equation, which is revisited in the next section.

### The Michaelis Menten model

Suppose now that the complex *C* is in a quasi-steady state [5]. This situation typically arises, for example, in cellular metabolic reactions when the substrate concentration is much higher than that of the enzyme, so that *C* quickly reaches equilibrium. In this regime, according to (8), the quasi-stationary concentration of *c* can be approximated by

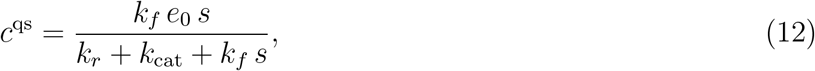

Replacing (12) into (7) yields the reduced dynamics for *s*, namely the standard MM equation

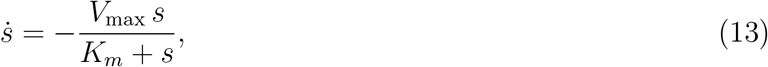

Where

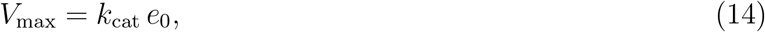

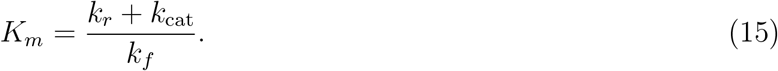

Equation (13) describes the substrate-to-product conversion through a single nonlinear ODE for *s*. If (12) holds, substitution into (6) and comparison with (13) shows that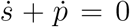, implying *p* = *h* − *s*, with *h* a positive constant determined by the initial conditions. The reaction rate *v* is defined as the right-hand side of (13) with positive sign, leading to the MM rate law [10]. Here, *V*_max_ denotes the limiting (or maximal) rate, reached as *s* → ∞, while *K*_*m*_ is the Michaelis (or half-saturating) constant, i.e. the substrate concentration at which the rate is *V*_max_*/*2 [8]. The parameters *V*_max_ (or *k*_cat_) and *K*_*m*_ are referred to as the MM parameters [3], and (15) is regarded as the constraint equation.

The solution *s*^*a*^(*t*) of Eq. (13) satisfying *s*^*a*^(0) = *s*_0_ provides an approximation of the solution *s*(*t*) of the original problem: as a consequence, an approximate value of *c*(*t*) is obtained by replacing *s*(*τ*) with *s*^*a*^(*τ*) in (10).

### Formulation of the inverse problem

Suppose that *e*_0_ is given. Equations (14) and (15) provide the MM parameters *V*_max_ and *K*_*m*_ in terms of the microscopic rates *k*_*f*_, *k*_*r*_, and *k*_cat_. Conversely: the rates *k*_*f*_, *k*_*r*_, and *k*_cat_ cannot be determined algebraically from *V*_max_ and *K*_*m*_, since only two equations are available for three unknowns. In practice, *k*_cat_ is directly related to *V*_max_ through *V*_max_ = *k*_cat_*e*_0_, so from now on *k*_cat_ is treated as a MM parameter (essentially equivalent to *V*_max_), and the actual unknowns are *k*_*f*_ and *k*_*r*_.

The analysis above shows that *k*_*f*_ and *k*_*r*_ must satisfy both (11) and the constraint (15): this system provides the starting point for estimating the unknown microscopic rates from parameters *K*_*M*_ and *k*_cat_. Rewriting (15) gives the equivalent identity

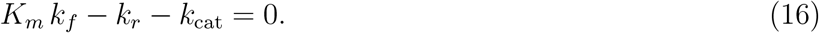

As for (11), it involves the substrate concentration *s*, which is unknown, but can be estimated as the solution *s*^*a*^ of the standard MM equation (13).

Replacing *s*^*a*^ into (11) and using (10) to obtain *c*^*a*^ = *c*[*s*^*a*^] yields the fundamental equation for *k*_*f*_ and

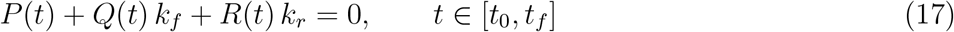

With

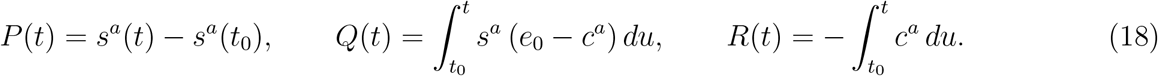

Here, [*t*_0_, *t*_*f*_] denotes the time window used for inversion. The rates *k*_*f*_ and *k*_*r*_ are then estimated by applying a numerical optimization method to the resolvent system formed by the constraint (16) and the fundamental equation (17).

### Inversion procedure

In the present approach, we solve a set of inverse problems for *k*_*f*_, *k*_*r*_, at fixed *K*_*m*_ and *k*_cat_; each problem corresponds to a different choice of the initial values *s*_0_ and *e*_0_. In order to avoid bias from the dependence of the solutions on these initial conditions, we define the final estimates of the microscopic rate constants as the geometric mean of the corresponding solutions. The solution procedure is structured into three main steps, summarized as follows.

Step 1. Under the assumption that *K*_*m*_ and *k*_cat_ are given, the two intervals ℐ_*s*_ and ℐ_*e*_ are defined, from which the respective initial values *s*_0_ and *e*_0_ are drawn. For any pair *s*_0_ ∈ ℐ_*s*_, *e*_0_ ∈ ℐ_*e*_ the solution of the MM equation is evaluated, providing a specific time interval [*t*_min_, *t*_max_] within which the inverse problem is solved.

Step 2. The rates *k*_*f*_ and *k*_*r*_ are estimated, based on (i) the given values of *K*_*m*_, *k*_cat_, *s*_0_, *e*_0_; (ii) the time interval [*t*_min_, *t*_max_]; (iii) the fundamental equation; (iv) the constraint equation. This constitutes the core of the inversion procedure.

Step 3. The geometric mean of the estimates of *k*_*f*_ and *k*_*r*_, that have been generated by varying *s*_0_ and *e*_0_, provides the required values of the microscopic rates.

Let us now comment on these three steps.

#### Step 1

We assume that *K*_*m*_ and *k*_cat_ are given. The initial values *s*_0_ and *e*_0_ are sampled respectively from the intervals ℐ_*s*_ = [*K*_*m*_*/*5, 1.5 *K*_*m*_], and ℐ_*e*_ = [*K*_*m*_*/*50, 0.5 *K*_*m*_], that have been chosen considering both the conditions for the application of the standard Quasi Steady-State approximation (QSSA) [10, 21]. The definitions take into account the fact that *K*_*m*_, *s*_0_, and *e*_0_ share the same concentration units. The bounds are motivated by simulation results and values reported in the literature. On average, the initial enzyme concentration is lower than that of the substrate, which is consistent with the known requirement for the validity of the MM model [22].

Next, we consider a pair *s*_0_ ∈ ℐ_*s*_, *e*_0_ ∈ ℐ_*e*_, and the time interval [0, *t*_*f*_], with *t*_*f*_ = 1500 s based on reconstruction results. The concentration *s*(*t*) is approximated on [0, *t*_*f*_] by the solution *s*^*a*^(*t*) of the MM equation (13), with *s*^*a*^(0) = *s*(0) = *s*_0_; moreover, the approximation of *c*(*t*) is given by *c*^*a*^(*t*) from (10), replacing *s* with *s*^*a*^. Simulations (see Figure 4, and also [23]) show that *c*^*a*^(*t*) rapidly rises, reaches a maximum, and then monotonically decays to 0 as *t* tends to infinity. Since the MM equation is derived under the quasi-stationary assumption, it seems reasonable to restrict attention to time instants *t*_min_, *t*_max_ within the interval where the system is close to a steady state, i.e. when the derivative 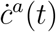is sufficiently small. The reaction is considered complete when only a small percentage *γ* ∈ (0, 30] of the initial substrate remains free. This leads to the definitions:

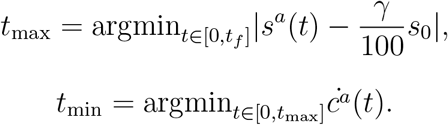

Following [10], we have set *γ* = 10.

Note that in our formulation the MM parameters are taken as (*K*_*m*_, *k*_cat_) rather than (*K*_*m*_, *V*_max_). Since *V*_max_ = *k*_cat_*e*_0_, varying the initial enzyme concentration *e*_0_ directly modifies *V*_max_. The case where *V*_max_ is given can thus be reduced to the present setting.

#### Step 2

The approximations *s*^*a*^(*t*) and *c*^*a*^(*t*), restricted to [*t*_min_, *t*_max_], are now used to estimate the unknown microscopic rates *k*_*f*_ and *k*_*r*_.

First, the time domain [*t*_min_, *t*_max_] is discretized by the partition *{t*_0_ = *t*_min_, *t*_1_, …, *t*_*n*_ = *t*_max_*}*. Evaluating (17) at each *t*_*i*_ leads to the system

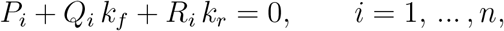

Where

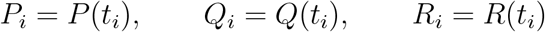

(see definitions in (18)).

To simplify the evaluation of the explicit expressions of the coefficients, we refer to the definitions (18) to observe that

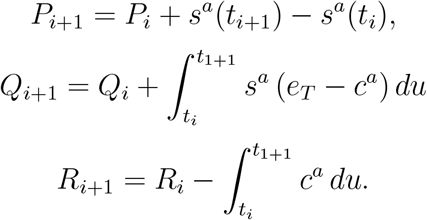

To identify (*k*_*f*_, *k*_*r*_), the penalized least-squares functional to be minimized, which incorporates the constraint (16) on the parameters, is defined as

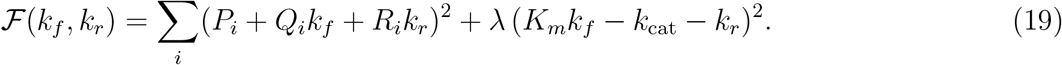

where the *L*^2^ norm has been considered and the (real, positive) factor *λ >* 0 controls the weight of the constraint. Expanding (19) yields the quadratic form

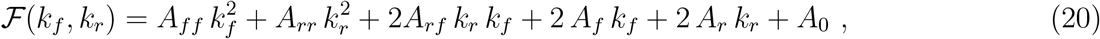

Where

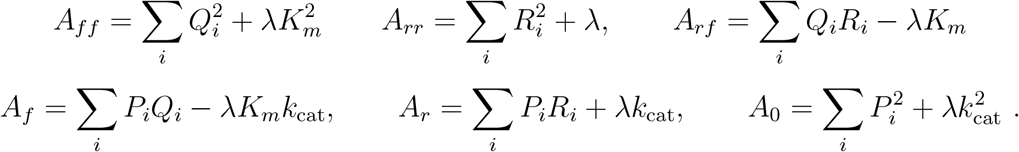

The stationarity condition leads to the linear system

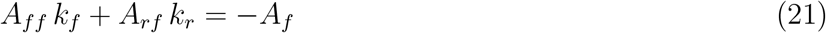

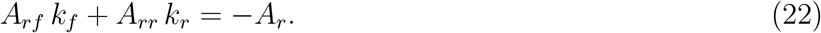

If the determinant of the coefficient matrix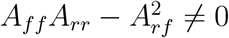, the solution is unique and given by

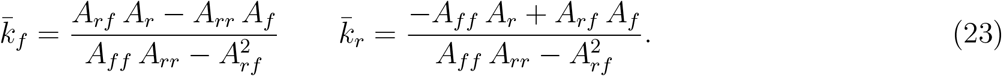

A proof that 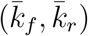 is the global minimizer of, provided that vectors **Q** and **R** are not both null, is given in the Supplementary Information.

Finally, to guarantee biologically meaningful estimates, both *k*_*f*_ and *k*_*r*_ are constrained to be non-negative. Therefore, whenever at least one negative result is encountered, the minimizers are instead computed numerically using MATLAB^®^ function fmincon, enforcing positivity constraints.

#### Step 3

First, a set of *n*_*p*_ ∈ ℕ pairs of initial concentrations *s*_0_ ∈ ℐ_*s*_ and *e*_0_ ∈ ℐ_*e*_ is sampled using the Latin Hypercube Sampling method [24]. For each sampled pair and fixed the MM parameters, the reconstruction procedure described in Step 2 is applied, yielding *n*_*p*_ estimates of (*k*_*f*_, *k*_*r*_). The final reconstructed values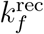 and 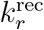are then defined as the geometric means of the *n*_*p*_ estimates.

This strategy minimizes dependency on specific initial conditions, and ensures robustness of the reconstruction, while keeping computational feasibility. In particular, restricting the sampling intervals and using the geometric mean mitigate the uncertainty caused by the low sensitivity of *k*_*r*_, which will be explored in the following section.

The overall procedure, named the Michaelis-Menten to Microscopic rate constants (MM2M) algorithm, is schematized in Algorithm (1) reported in the Supplementary material.

## Results

### Sensitivity analysis

We first applied sensitivity analysis to investigate how *s* depends on rates *k*_*f*_ and *k*_*r*_ [15, 25, 26, 27, 28, 29]. Specifically, we compared substrate concentration curves obtained by solving the direct problem for different parameter values. The results provide guidance for formulating and addressing the inverse problem.

Here, we present numerical experiments illustrating how the substrate concentration *s(t*) evolves when the dissociation and association microscopic rates are varied independently. To highlight the relative influence of *k*_*f*_ and *k*_*r*_, we kept the catalytic rate fixed at *k*_cat_ = 1 s^−1^ and compared two representative scenarios: (i) *k*_*f*_ = 0.500 nM^−1^s^−1^, *k*_*r*_ = 0.005 s^−1^, (ii) *k*_*f*_ = 0.005 nM^−1^s^−1^, *k*_*r*_ = 0.500 s^−1^. These two parameter sets essentially invert the relative magnitude of *k*_*f*_ and *k*_*r*_, providing a balanced framework to test whether the sensitivity of *s*(*t*) to the dissociation constant changes when the ratio *k*_*f*_ */k*_*r*_ is reversed. Figure 1 reports the resulting time courses of *s* obtained from system (7) -(8). In panels (A) and (C), *k*_*r*_ is fixed, while *k*_*f*_ varies over a set of 21 logarithmically spaced values within the range [10^−4^, 1] nM^−1^s^−1^. In panels (B) and (D), The roles of *k*_*f*_ and *k*_*r*_ are reversed. In all cases, the initial conditions are set to *s*(0) = 20 nM, *c*(0) = 0 nM, *e*(0) = 10 nM.

**Figure 1.**
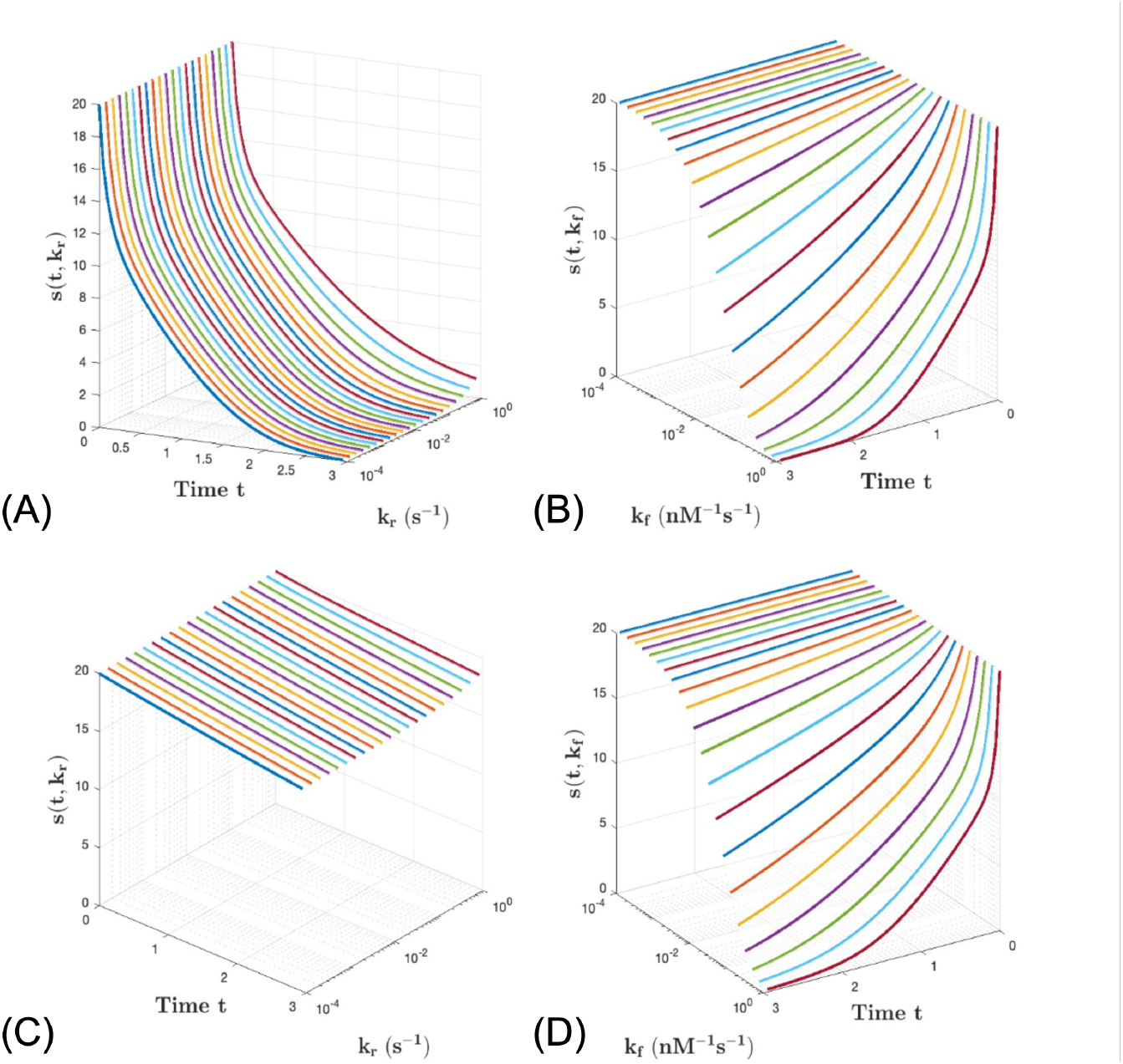
Behavior of the concentration of the substrate *s* over time. In each panel, *k*_cat_ = 1 s^−1^, *s*_0_ = 20 nM and *e*_0_ = 10 nM, while either *k*_*f*_ or *k*_*r*_ is fixed: (A) *k*_*f*_ = 0.500nM^−1^s^−1^; (B) *k*_*r*_ = 0.005 s^−1^; (C) *k*_*f*_ = 0.005 nM^−1^s^−1^; (D) *k*_*r*_ = 0.500s^−1^ is fixed. When not fixed, *k*_*r*_ always varies logarithmically over 21 values from 10^−4^ to 1s^−1^, and similarly *k*_*f*_ spans the range from 10^−4^ to 1 nM^−1^ s^−1^).

In both couples of plots, *s* appears significantly more influenced by *k*_*f*_ than by *k*_*r*_, a pattern consistently observed in multiple cases. Indeed, in both scenarios, the curves dependent on *k*_*r*_ are nearly undistinguishable, especially for the smaller values of *k*_*r*_. This suggests a higher sensitivity of *s* to the association constant and a lower sensitivity to the dissociation one. Note that, even inverting the ratio between the two rate constants, the low sensitivity of *s* to *k*_*r*_ variations remains nearly unchanged. Furthermore, a strong similarity can be appreciated between panels (B) and (D), where *k*_*f*_ assumes the same set of values and setting *k*_*r*_ = 0.500 s^−1^ or *k*_*r*_ = 0.005 s^−1^ has no apparent impact on the substrate’s behavior.

The sensitivity analysis was then enriched by observing the local sensitivities of *s* and *c* with respect to *k*_*f*_ and *k*_*r*_, defined by the time course of the derivatives

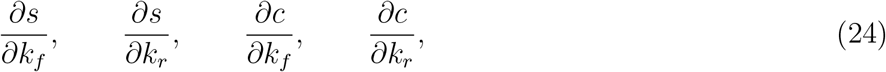

which were evaluated along any solution *s*(*t*), *c*(*t*) of the system (7), (8), with *k*_*f*_ and *k*_*r*_ fixed. These sensitivity functions were computed by solving the corresponding system of four coupled ODEs (details are in the Supplementary Information) in the same two scenarios used for the previous analysis. The results, shown in Figure 2, confirm the predominant influence of the association rate: in most cases, the magnitude of ∂*s/*∂*k*_*f*_ dominates over that of ∂*s/*∂*k*_*r*_. Only in a limited portion of the time course ∂*s/*∂*k*_*r*_ briefly exceeds ∂*s/*∂*k*_*f*_, suggesting that in certain cases, likely when *k*_*r*_ is relatively high, even small variations in *k*_*r*_ may still produce a detectable effect on the substrate dynamics, making it possible to recover at least its order of magnitude. Further discussion of this aspect will be provided in the numerical results section.

**Figure 2.**
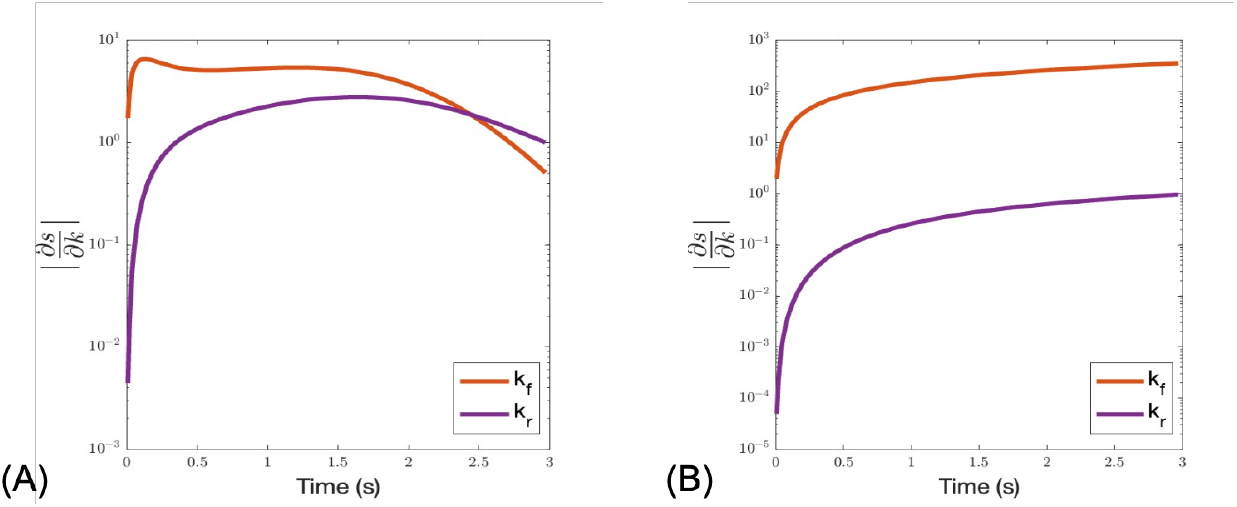
Derivatives of the function *s*, describing the dynamics of the substrate concentration, with respect to *k*_*f*_ and *k*_*r*_ in cases (A) *k*_*f*_ = 0.500 nM^−1^s^−1^, *k*_*r*_ = 0.005 s^−1^, *k*_cat_ = 1 s^−1^, and (B) *k*_*f*_ = 0.005 nM^−1^s^−1^, *k*_*r*_ = 0.500 s^−1^, *k*_cat_ = 1 s^−1^. In both panels, *s*_0_ = 20 nM and *e*_0_ = 10 nM.

While these are just representative examples, the observed trends suggest that reconstructing *k*_*r*_ is likely to be significantly more error-prone than estimating *k*_*f*_. Therefore, the assignment of a precise numerical value to *k*_*r*_, as a result of the inversion procedure, should be done with caution.

### Numerical validation of the inversion technique

A number of numerical experiments have been considered to ascertain the reliability of the proposed approach for converting the MM constants into the corresponding microscopic rate constants. Each simulation has been developed according to the following scheme: (i) assignment of the ground values to *k*_*f*_, *k*_*r*_ and *k*_cat_; (ii) evaluation of the corresponding MM parameter *K*_*M*_ ; (iii) reconstruction of the values of the microscopic association and dissociation rates from *K*_*M*_ and *k*_cat_ by application of the inversion procedure. The values of the microscopic rate constants have been sampled following a logarithmic distribution across specific intervals for each constant: *k*_*f*_ ∈ [10^−4^, 1] nM^−1^s^−1^, *k*_*r*_ ∈ [10^−3^, 3] s^−1^, *k*_cat_ ∈ [10^−2^, 5] s^−1^. These intervals were inspired by the trends of the values reported in the literature for reactions defining different pathways (see [30, 31]).

For each reconstruction associated with a given triple *k*_*f*_, *k*_*r*_, and *k*_cat_, *n*_*p*_ = 10 simulations have been performed, with different values for the initial concentrations of substrate and enzyme, selected as explained in the section discussing the inversion procedure.

As far as time is concerned, as already mentioned, we have set *t*_*f*_ = 1500 s. The interval [*t*_min_, *t*_max_] has been found following the prescriptions of the previous section, where the value of *γ* has been set to 10. An equispaced discretization of [*t*_min_, *t*_max_] with a time step of 1*/*100 s defines the vector **t**. The parameter *λ* has always been considered as equal to 1 so as to give the same weight to the two penalty functions that compose ℱ (see (19)).

Table 1 presents the results of a single simulation, with the values of the ground rate constants fixed at *k*_*f*_ = 0.0006 nM^−1^ s^−1^, *k*_*r*_ = 1.8453 s^−1^, and *k*_cat_ = 4.1386 s^−1^. The geometric mean of the reconstructed constants is shown in the last row of the table. The last two columns display the ratios between the reconstructed values of *k*_*f*_ and *k*_*r*_ and their respective ground truth values. The results obtained for *k*_*f*_ align well with expectations and, in fact, are rather close to the true ones. In this specific case, the results highlight that averaging over different initial concentration values of the species is an effective strategy to obtain a ratio 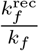 that is closer to 1, compared to those obtained from individual simulations, with *s*_0_ and *c*_0_ fixed. As expected, the reconstructions of *k*_*r*_ are more subject to error than those of *k*_*f*_. Also in this case, the geometric mean provides a more reliable value of *k*_*r*_ as it effectively averages across the orders of magnitude of the reconstructed values, without being overly influenced by individual deviations.

**Table 1.**
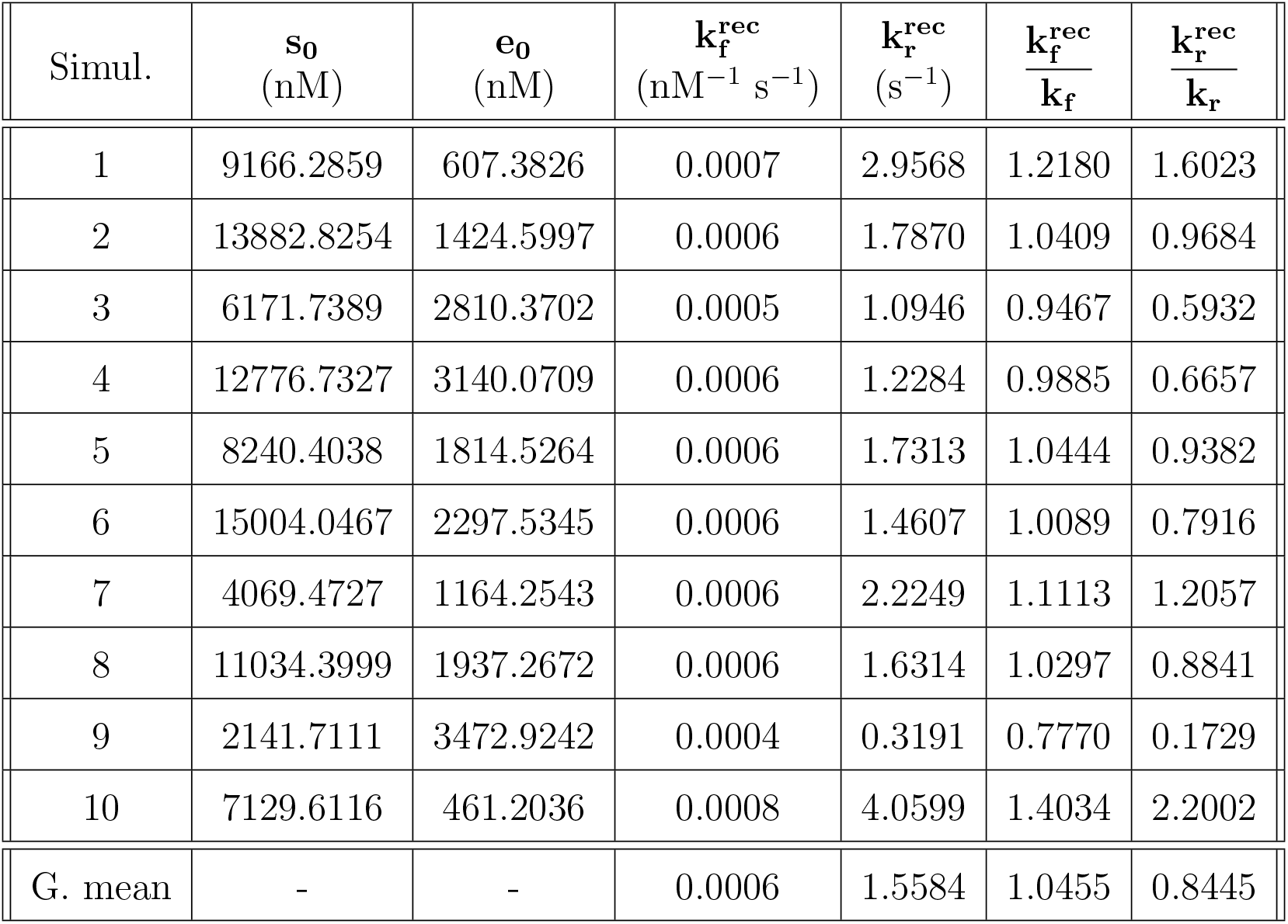
Ten simulations of rate conversion, from MM format to microscopic rate constants. The rates are fixed as: *k*_*f*_ = 0.0006 nM^−1^ s^−1^, *k*_*r*_ = 1.8453 s^−1^, *K*_*M*_ = 10551.70 nM and *k*_cat_ = 4.1386 s^−1^, but the values for *s*_0_ and *e*_0_ are different in each simulation. Columns **s**_**0**_ and **e**_**0**_ show the fixed values for substrate and enzyme initial concentrations, 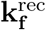 and 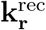 the reconstructed microscopic rates, and the last two columns present the ratio between reconstructed values and the ground truth. The last row displays the results obtained by averaging (geometric mean) the rates *k*_*f*_ and *k*_*r*_ from the previous rows across different initial concentrations of enzyme and substrate.

| Simul. | $s_0$<br>(nM) | $e_0$<br>(nM) | $k_f^{\text{rec}}$<br>(nM<sup>-1</sup> s<sup>-1</sup>) | $k_r^{\text{rec}}$<br>(s<sup>-1</sup>) | $\frac{k_f^{\text{rec}}}{k_f}$ | $\frac{k_r^{\text{rec}}}{k_r}$ |
| --- | --- | --- | --- | --- | --- | --- |
| 1 | 9166.2859 | 607.3826 | 0.0007 | 2.9568 | 1.2180 | 1.6023 |
| 2 | 13882.8254 | 1424.5997 | 0.0006 | 1.7870 | 1.0409 | 0.9684 |
| 3 | 6171.7389 | 2810.3702 | 0.0005 | 1.0946 | 0.9467 | 0.5932 |
| 4 | 12776.7327 | 3140.0709 | 0.0006 | 1.2284 | 0.9885 | 0.6657 |
| 5 | 8240.4038 | 1814.5264 | 0.0006 | 1.7313 | 1.0444 | 0.9382 |
| 6 | 15004.0467 | 2297.5345 | 0.0006 | 1.4607 | 1.0089 | 0.7916 |
| 7 | 4069.4727 | 1164.2543 | 0.0006 | 2.2249 | 1.1113 | 1.2057 |
| 8 | 11034.3999 | 1937.2672 | 0.0006 | 1.6314 | 1.0297 | 0.8841 |
| 9 | 2141.7111 | 3472.9242 | 0.0004 | 0.3191 | 0.7770 | 0.1729 |
| 10 | 7129.6116 | 461.2036 | 0.0008 | 4.0599 | 1.4034 | 2.2002 |
| G. mean | - | - | 0.0006 | 1.5584 | 1.0455 | 0.8445 |

Table 2 reports 12 representative estimates of *k*_*f*_ and *k*_*r*_, selected from the total of 50 computed cases, together with the corresponding ratios 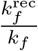 and 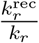. According to these reconstructions, the solution 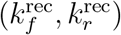 provides a fairly precise approximation of the true value of *k*_*f*_, which appears to be of the same order of magnitude. In contrast, the reconstruction of *k*_*r*_ is noticeably less accurate, as expected from the results of sensitivity analysis. This is illustrated in Figure 3, which presents the distribution of the ratios 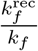 and 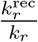 in the 50 considered scenarios. The plot highlights how the reconstructed forward rates closely match the original values, whereas the backward rates exhibit a significantly different behavior.

**Table 2.** Twelve reconstructions of microscopic rate constants, from MM constants. Columns **k**_**f**_ and **k**_**r**_ show the true values for these two rate constants, **K**_**M**_ and **k**_cat_ the corresponding Michaelis-Menten rates, 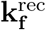 and 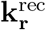 the reconstructed microscopic rates, and the last two columns present the ratio between reconstructed values and the ground truth.

| Simul. | $\mathbf{k}_f$<br>($\text{nM}^{-1} \text{s}^{-1}$) | $\mathbf{k}_r$<br>($\text{s}^{-1}$) | $\mathbf{K}_M$<br>($\text{nM}$) | $\mathbf{k}_{\text{cat}}$<br>($\text{s}^{-1}$) | $\mathbf{k}_f^{\text{rec}}$<br>($\text{nM}^{-1} \text{s}^{-1}$) | $\mathbf{k}_r^{\text{rec}}$<br>($\text{s}^{-1}$) | $\frac{\mathbf{k}_f^{\text{rec}}}{\mathbf{k}_f}$ | $\frac{\mathbf{k}_r^{\text{rec}}}{\mathbf{k}_r}$ |
| --- | --- | --- | --- | --- | --- | --- | --- | --- |
| 1 | 0.0008 | 0.4790 | 6869.66 | 4.8287 | 0.0011 | 2.0168 | 1.40 | 4.21 |
| 2 | 0.3040 | 0.0485 | 6.71 | 1.9917 | 0.3132 | 0.0032 | 1.03 | 0.07 |
| 3 | 0.0007 | 0.0013 | 22.71 | 0.0140 | 0.0009 | 0.0048 | 1.31 | 3.81 |
| 4 | 0.4725 | 0.0020 | 0.07 | 0.0327 | 0.5694 | 0.0058 | 1.21 | 2.89 |
| 5 | 0.0090 | 0.0024 | 18.78 | 0.1664 | 0.0129 | 0.0593 | 1.43 | 24.28 |
| 6 | 0.0280 | 0.0075 | 28.47 | 0.7893 | 0.0398 | 0.2708 | 1.42 | 36.23 |
| 7 | 0.1158 | 2.2653 | 45.01 | 2.9465 | 0.0824 | 0.1640 | 0.71 | 0.07 |
| 8 | 0.0118 | 0.1573 | 59.39 | 0.5464 | 0.0133 | 0.1921 | 1.13 | 1.22 |
| 9 | 0.0015 | 0.6912 | 3230.72 | 4.2804 | 0.0020 | 1.8305 | 1.33 | 2.65 |
| 10 | 0.0006 | 0.0930 | 7541.26 | 4.1564 | 0.0009 | 1.8267 | 1.62 | 19.65 |
| 11 | 0.0002 | 0.1619 | 3885.59 | 0.6555 | 0.0003 | 0.2680 | 1.21 | 1.66 |
| 12 | 0.0899 | 0.6654 | 52.73 | 4.0750 | 0.0882 | 0.0577 | 0.98 | 0.09 |

**Figure 3.**
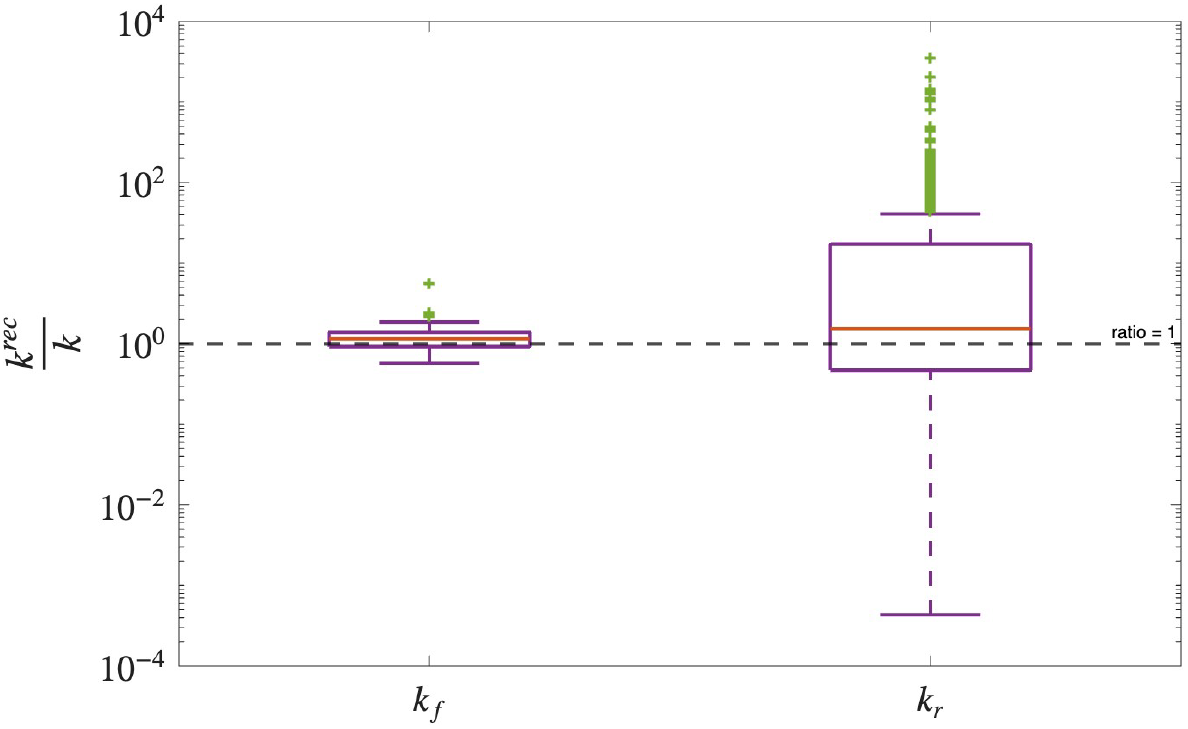
Ratios between rate constants 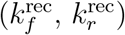 reconstructed by the MM2M algorithm and their true values (*k*_*f*_, *k*_*r*_). (A) Results from 500 single simulations (10 runs for each of the 50 reconstructions), before applying the geometric mean. (B) Results from the corresponding 50 reconstructions obtained after applying the geometric mean.

However, when 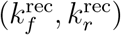 have been replaced into the system (7), (8) and the corresponding direct problem has been solved, the reconstructed curves *s*^rec^ and *c*^rec^ have closely approximated the true curves *s* and *c*, demonstrating the overall reliability of the approach. A similar result was shown in [10], in the reconstruction of the MM parameters *K*_*M*_ and *V*_max_ from time course experiments. Figure 4 provides a comparison between the true curves *c* and *s* (colored solid lines), their reconstructions *s*^*MM*^ and *c*^*MM*^ obtained via the MM equation (black solid lines), and their estimates *s*^rec^ and *c*^rec^ produced by our model (colored dashed lines). Note that *c*^*MM*^ is computed using equation (10), where *s* is replaced by its MM approximation *s*^*MM*^ from equation(13). Panels (A), (B), and (C) show the results obtained after reconstructing three different couples of association and dissociation rate constants, respectively, those related to reconstructions labeled by 12 (*k*_*f*_ = 0.0899 nM^−1^ s^−1^, *k*_*r*_ = 0.0076 s^−1^), 6 (*k*_*f*_ = 0.0280 nM^−1^ s^−1^, *k*_*r*_ = 0.0107 s^−1^), and 10 (*k*_*f*_ = 0.0006 nM^−1^ s^−1^, *k*_*r*_ = 0.0084 s^−1^). The order of the panels reflects both the accuracy achieved by our method and the limited performance of the MM reconstruction. Note that these aspects are influenced by the amplitude of the reconstruction time window, which is determined by the intrinsic speed of the reaction. The estimate derived from the MM equation becomes accurate once *c* enters a quasi-steady-state regime, in line with the underlying assumptions of the MM model. For example, the *s* curve reconstructed using the MM method and shown in panel (C) closely matches the true trajectory starting from approximately *t* = 400 s, which marks the onset of a quasi-steady state for the species *C*. Instead, in the early stages of the reaction the approximation produced by our algorithm yields noticeably higher accuracy. Furthermore, our algorithm generally outperforms the MM approach in reconstructing the *c* curve, as illustrated in panels (B) and (C). Panel (A) depicts a scenario in which the reaction takes longer to reach 90% completion, resulting in the three curves appearing almost identical; a closer inspection confirms they are very similar, though not perfectly overlapping. Note the different scale on both x and y axis, which vary depending on two factors: the values of *s*_0_ for each simulation -which have been chosen respectively as the initial condition for reconstruction 12 simulation 1, rec. 6 sim.9, and rec. 10 sim. 5-, and the time required for the substrate to deplete to 10% 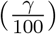 of its initial value, meaning the reaction is considered 90% complete.

One limitation of averaging results over the *n*_*p*_ runs, rather than relying on the individual reconstructions obtained for fixed *s*_0_ and *e*_0_, is that the constraint equation (16) is not guaranteed to be satisfied in general.

Since Table 2 shows that reconstructions labeled by 1 and 10 are remarkably precise not only in terms of *k*_*f*_, but also in *k*_*r*_, an unexpected outcome based on the observations provided in the Section concerned with sensitivity analysis, we focused on reconstruction n. 10 and therefore analyzed how the behavior of *s* changes when *k*_*f*_ is fixed at its ground value 0.0006 nM^−1^ s^−1^, while *k*_*r*_ varies on a logarithmic scale centered at its reference value 0.0084 s^−1^, and viceversa.

The *s* we aim to reconstruct with our method is shown in purple and is indicated by the red arrow in Figure 5. As illustrated in the plot, when *k*_*r*_ is close to 0.008 nM^−1^s^−1^, small variations of this parameter can lead to significant differences in the reconstruction of *s*, particularly when the approximate value for *k*_*r*_ is larger than its ground truth. This indicates that, in this specific case, the reconstruction is also sensitive to this parameter.

**Figure 4.**
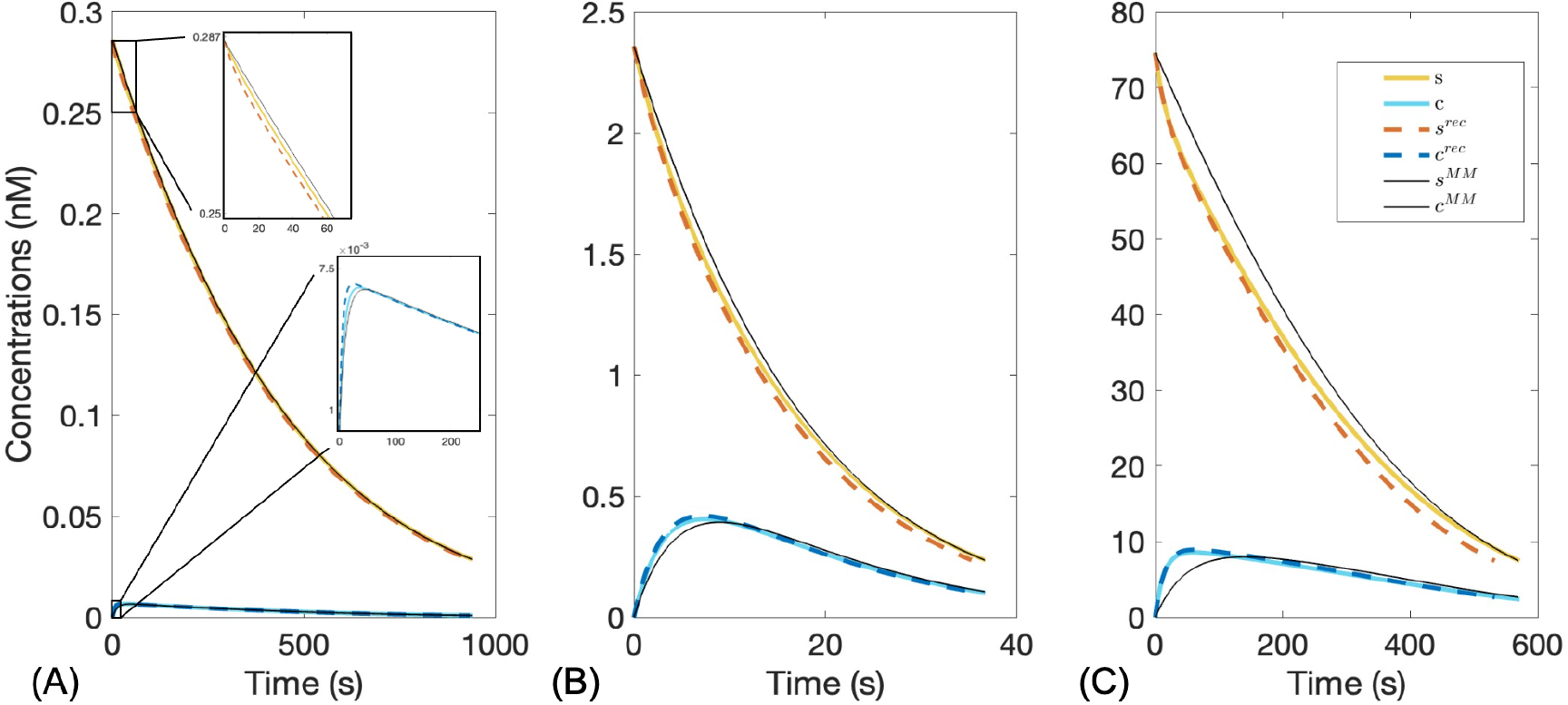
Comparison between three representations of the substrate and complex concentrations *s*(*t*) and *c*(*t*): the ground truth solutions (coloured solid lines), the approximations from the MM equation (black solid lines), and the estimates obtained with our algorithm (coloured dashed lines). Results associated respectively to reconstructions of Table 2 n. (A) 12 (*s*_0_ and *e*_0_ from simul. 1), (B) 6 (*s*_0_ and *e*_0_ from simul. 9), (C) 10 (*s*_0_ and *e*_0_ from simul. 5). In panel (A), a zoom-in of the initial behavior of *s* and *c* is provided. See the Supplementary Information for further information about the selected values for *s*_0_ and *e*_0_ in the single reconstructions.

**Figure 5.**
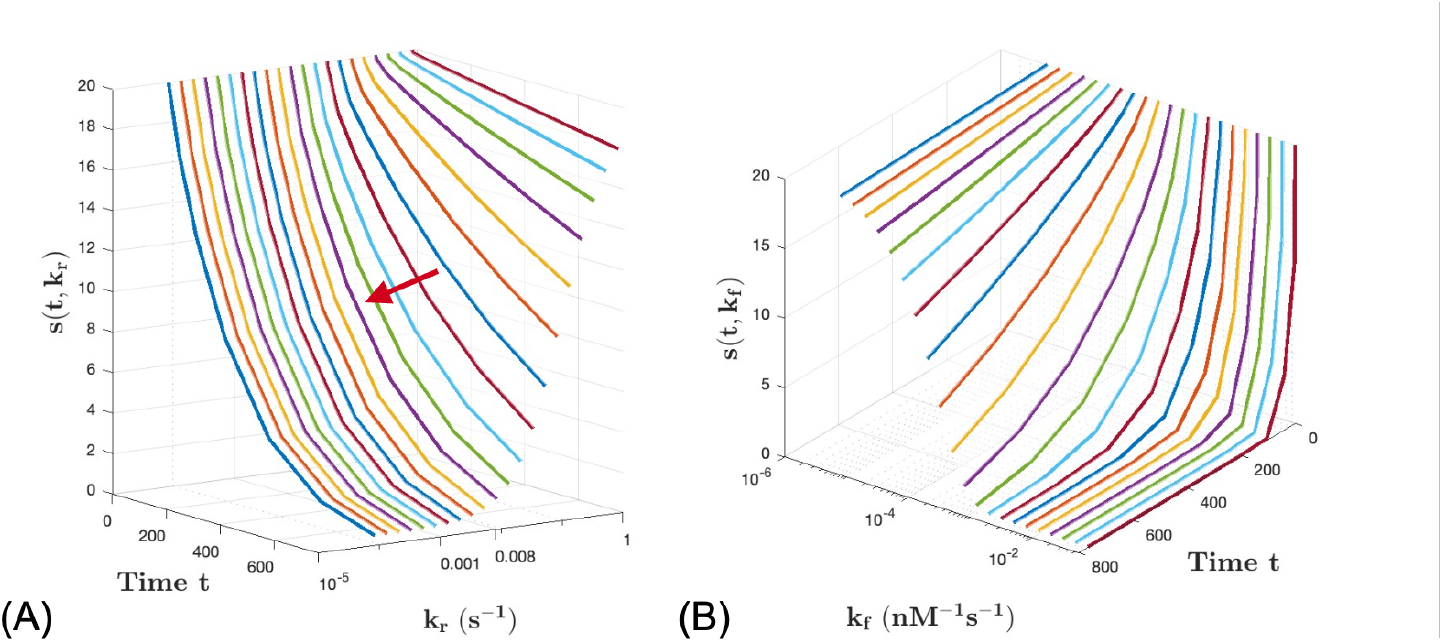
Behavior of the concentration *s* over time. Panel (A): *k*_*f*_ = 0.0006nM^−1^s^−1^ is fixed, while *k*_*r*_ takes 21 values, scaled logarithmically from 0.0084s^−1^, ground truth of *k*_*r*_ for case n. 5. Panel (B): *k*_*r*_ = 0.0084s^−1^ is fixed, while *k*_*f*_ takes 21 values, scaled logarithmically from 0.0006nM^−1^s^−1^, ground truth of *k*_*f*_ for case n. 10.

### Case study on reactions within the PI3K/AKT/mTOR pathway

A second numerical simulation was performed using literature-derived values of MM parameters (*K*_*M*_ and *k*_*cat*_) associated with specific reactions of the PI3K/AKT/mTOR pathway [32, 33, 34]. This signaling cascade plays a pivotal role in both neurodegenerative diseases and cancer as well as in metabolic disorders such as diabetes and obesity, making it one of the most interesting signaling pathways, thoroughly studied as therapeutic target. Indeed, it regulates fundamental cellular processes such as growth, metabolism, and protein translation.

For a subset of enzyme-catalyzed reactions within this pathway, the literature reports only MM parameters, while the corresponding microscopic rate constants remain unavailable. Ten of these reactions were therefore selected for our study, with the aim of reconstructing the underlying microscopic rate constants through the MM2M algorithm in the absence of direct experimental measurements. The detailed list of the selected reactions is reported in Table 3. The parameters for the algorithm have been selected as in the previous set of simulations: *t*_*f*_ = 1500 s, *γ* = 10, and *λ* = 1. The time vector **t** spans the interval [*t*_min_, *t*_max_] uniformly discretized with a time step of 1*/*100 s. Results are summarized in Table 4, which presents, for each reaction, the available MM parameters together with the reconstructed microscopic rates.

**Table 3.**
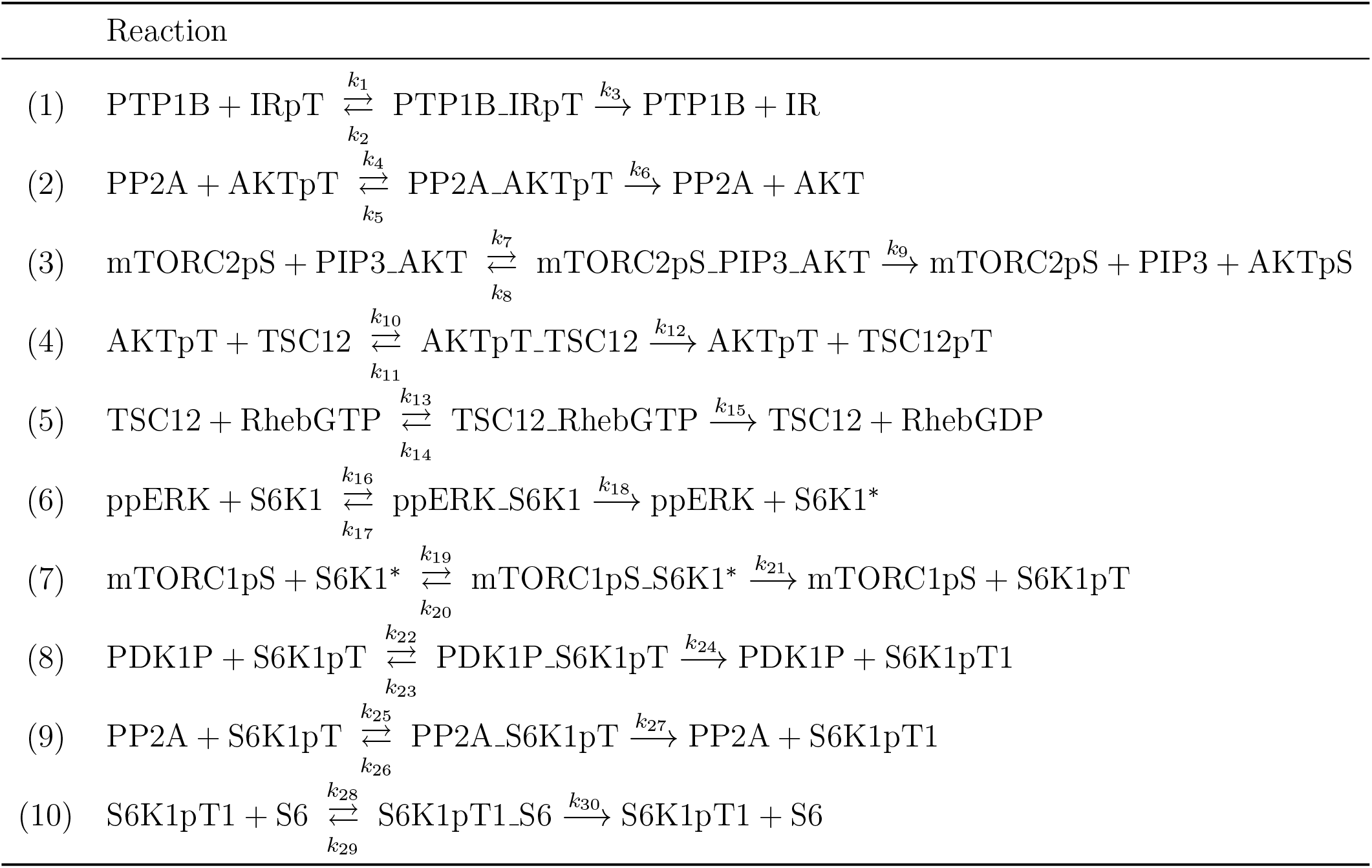
List of enzyme-catalyzed reactions selected from the PI3K/AKT/mTOR pathway.

**Table 4.**

| Reaction | $K_m$ (nM) | $k_{cat}$ (s<sup>-1</sup>) | $k_f^{rec}$ (nM<sup>-1</sup>s<sup>-1</sup>) | $k_r^{rec}$ (s<sup>-1</sup>) | Reference |
| --- | --- | --- | --- | --- | --- |
| (1) | 7500 | 11.3 | 0.0015 | 0.0010 | [35] |
| (2) | 48000 | 1.8 | 0.0012 | 0.4842 | [36] |
| (3) | 800 | 20 | 0.0252 | 0.0077 | [36] |
| (4) | 10300 | 6 | 0.0007 | 0.0027 | [36] |
| (5) | 300 | 20 | 0.0672 | 0.0024 | [36] |
| (6) | 17300 | 1 | 0.0004 | 5.767 | [36] |
| (7) | 800 | 0.06 | 0.0001 | 0.0563 | [36] |
| (8) | 10000 | 1 | 0.0007 | 1.5994 | [36] |
| (9) | 8800 | 1 | 0.0009 | 6.343 | [36] |
| (10) | 250 | 0.01 | 0.00007 | 0.0073 | [36] |

## Discussion

### Aim

In this work, we have revisited the classical model of enzymatic reactions (3)-(6), consisting of a system of four ODEs describing the evolution of four species concentrations, along with its well-known simplification given by the MM standard equation (13). The ODE system depends on the three rate constants *k*_*f*_, *k*_*r*_, and *k*_cat_, while the MM equation involves the parameters *K*_*m*_ and *V*_max_ (or *k*_cat_ = *V*_max_*/e*_0_, since *e*_0_ is considered known). Our specific aim has been to estimate the microscopic reaction rate constants, *k*_*f*_ and *k*_*r*_, in terms of the MM parameters, *K*_*m*_ and *k*_cat_. To this end, we have formulated and solved a family of inverse problems.

The main motivation has come from the observation that many experimental and computational methods available for the estimate of the MM parameters exist, and that the resulting values may be found in the literature in many cases of interest. In contrast, the underlying rate constants are generally more difficult to access, despite their crucial role in providing specific information on each reaction and their necessity in formulating reaction networks based on the mass-action law. In this regard, we recall that a network characterized by chemical reactions obeying the mass-action law is provided with useful mathematical properties, coming from the polynomial form of the system of ODEs, that enable the application of powerful analytical and computational tools for the analysis of reaction networks [15, 16, 17, 37]. Beyond these formal advantages, chemical reaction networks constitute a very versatile framework for the simulation of the dynamics of the chemical species within a given pathway or cellular context. A computational study of these networks allows researchers to observe their behaviour under physiological, perturbed, or drug-treated conditions. Therefore, constructing such networks - which are often not readily available in literature - is essential for investigating the pathways they encode and for uncovering the mechanisms driving processes such as tumor development and progression [19], as well as other pathological conditions.

### Intrinsic limitations of the results

Under the assumption that the data *K*_*m*_ and *k*_cat_ are known, the algorithm proposed in this work provides estimates of the corresponding microscopic rate constants *k*_*f*_ and *k*_*r*_. In our opinion, these results should be interpreted primarily as indicative of the expect orders of magnitude of these quantities, rather than as accurate values in specific biological systems. Indeed, the pertinent parameters for an in vivo enzymatic reaction are influenced by a moltitude of factors, including the cellular context, the possible existence of several substrates, the complex multiple molecular interactions, the uncertainties in the estimates of the MM parameters, and in the inversion algorithm itself [3]. An effective account of the consequences of these features can only be achieved through a fine tuning of the preliminary estimates generated by the standard inversion procedure. In other words, a reliable practical application of a chemical reaction network model may require a further improvement of the estimate of the parameter values, based on an appropriate, dedicated set of experimental measurements.

Furthermore, the choice of initial substrate and enzyme concentrations was constrained by the requirements of the sQSSA. Eventually, approximating *s* through the total Quasi Steady-State Approximation (tQSSA) [38, 39, 40] would relax this limitation, since it remains valid even when substrate and enzyme concentrations are comparable. However, extending the method to the tQSSA framework is nontrivial, as the resulting equations are more complex. For this reason, our algorithm is best regarded as a first step, tailored to regimes where the sQSSA holds, with possible generalizations left for future work.

### Limitations from sensitivity analysis

The unknown rates have been estimated by solving an inverse problem, but this approach relies on the satisfaction of certain conditions. A preliminary analysis of solutions to the associated direct problems has shown that the substrate concentration *s* is highly sensitive to changes in the forward rate constant *k*_*f*_, but only weakly dependent on *k*_*r*_. This implies, in particular, that variations in the value of *k*_*f*_ induce modifications of the time curve of *s*, whereas it is only weakly altered by changes in *k*_*r*_. This behaviour has been derived by applying both global and local sensitivity analysis techniques [26, 27, 28], in order to investigate the sensitivity of the simple model of enzymatic reactions to the rates *k*_*f*_ and *k*_*r*_. As expected, preliminary numerical reconstructions through the inversion procedure have shown that *k*_*f*_ can be estimated with reasonably accuracy from the approximated time profile of *s*, whereas the estimation of *k*_*r*_ is more prone to significant error. Accordingly, reducing the arbitrariness in the determination of *k*_*r*_ may be a further goal of the inversion procedure.

### Sources of error

We can identify at least three error sources. The first concerns the imprecise value of the parameters *K*_*m*_ and *k*_cat_, which have been regarded as exact despite being estimated through a complex combination of experimental and computational methods. A second source of error arises from replacing the ground truth *s*(*t*) in the fundamental equation with an estimate *s*^*a*^(*t*) obtained by applying the MM approximation (i.e. the sQSSA) is another source of error. A third potential source of inaccuracy comes from the numerical integration of the MM equation itself.

Despite these limitations, the overall impact of these errors does not appear to be sufficient to compromise the reconstruction of the time courses of *s*(*t*) and *c*(*t*) derived from the reconstructed parameters. A similar behavior can be observed in the work of Stroberg and Schnell [10], where accurate reconstructions of the species dynamics were obtained even when the estimated parameter values showed significant deviations from the true ones. This study also highlighted how selecting an initial substrate concentration close to *K*_*m*_, along with a significantly lower initial enzyme concentration, improves the accuracy of the reconstructed time courses — a strategy we have likewise implemented in our analysis.

### Variants

(i) The inversion procedure remains applicable even when the catalytic constant *k*_cat_ and the time course *s*^exp^ of the substrate concentration over the time interval 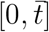 are available - for instance, through direct measurements. In this case, *s* is replaced by *s*^exp^ in the expression (10) of *c*, which is then substituted, along with *s*, into identity (11), giving the fundamental equation. The inverse problem is then solved over the time interval 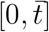, using the cost function defined in (19) without the constraint penalty term. This yields a rough estimate of *k*_*f*_ and *k*_*r*_. Subsequently, *K*_*m*_ is defined as *K*_*m*_ = (*k*_*r*_ + *k*_cat_)*/k*_*f*_. (ii) Another different scenario could arise: suppose that *K*_*m*_ is still known, but instead of *k*_cat_ the other input parameter is *V*_max_. In this case, the two intervals ℐ_*s*_ and ℐ_*e*_ can still be introduced, as in Sect., since their definitions depend only on *K*_*m*_. Then, at each *j*-th iteration a different pair (*s*_0_, *e*_0_)_*j*_ ∈ ℐ_*s*_ *×* ℐ_*e*_ is selected according to the previously described procedure. The value of *k*_*cat*_ is deduced straightforwardly as *k*_cat_ = *V*_max_*/e*_0_, so that the simulation of the corresponding system can proceed. After a predefined number of iterations, the geometric mean is applied to the estimated rates, yielding the final reconstructions 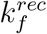and 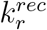.

## Conclusions

In this study, an effective method was developed and analyzed for estimating the microscopic kinetic parameters *k*_*f*_ and *k*_*r*_ starting from the experimentally measurable MM parameters *K*_*m*_ and *k*_cat_. Through an inversion approach based on a new identity for the substrate concentration and the subsequent replacement of *s* with its approximation given by the solution of the MM equation, a good estimate for the coefficients of the system of ODEs that models the enzymatic reaction has been reached. Numerical simulations realized under various initial conditions show that the reconstruction of *k*_*f*_ is generally accurate and stable, compared to *k*_*r*_ estimation, which is more prone to errors, confirming the sensitivity analysis of the model.

Nevertheless, the validity of the approach is supported by the good agreement between the reconstructed and true concentration time courses of substrate and enzyme-substrate complex. Ultimately, this work not only contributes to the quantitative understanding and analysis of enzymatic kinetics at a microscopic level, but also allows the application of mass-action kinetics to mathematically model large-scale biochemical systems via systems of ordinary differential equations.

## Author Contributions

GC and SB jointly conceived the method and the mathematical modeling. SB planned and performed the computational experiments, and validated the results. GC and SB wrote the text of the manuscript. All authors revised and approved the final version.

## Acknowledgements

SB, SS, and MP were partially found by Hub Life Science - Digital Health (LSH-DH) PNC-E3-2022-23683267 - Progetto DHEAL-COM - CUP: D33C22001980001, founded by Ministero della Salute within “Piano Nazionale Complementare al PNRR Ecosistema Innovativo della Salute - Codice univoco investimento: PNC-E.3”.

## Abbreviations

*C*: enzyme-substrate complex
*c*: complex concentration
*c*_0_: complex initial concentration
CRN: Chemical Reaction Network
*E*: enzyme
*e*: enzyme concentration
*e*_0_: enzyme initial concentration
*k*_**cat**_: microscopic catalitic constant
*k*_*f*_: microscopic forward constant
*K*_*m*_: Michaelis–Menten constant
*k*_*r*_: microscopic reverse constant
MM: Michaelis–Menten
ODE: Ordinary Differential Equation
*P*: product
*p*: product concentration
*S*: substrate
*s*: substrate concentration
*e*_0_: substrate initial concentration
*V*_**max**_: catalytic maximum velocity

