## Supplementary Information for "From Michaelis–Menten parameters to microscopic rate constants: an inversion approach for enzyme kinetics"

### SI1. The MM2M algorithm.

In this section, we provide a formal and detailed description of the MM2M algorithm introduced and analyzed in the manuscript.

---

**Algorithm 1:** The MM2M algorithm

---

**Input** : $K_M, k_{\text{cat}}, t_f, \lambda \in (0, +\infty)$, $\gamma \in (0, 30]$, $n_p \in \mathbb{N}$
**Output:** Estimated $k_f^{\text{rec}}$ and $k_r^{\text{rec}}$
Sample $n_p$ pairs $\{(e_0^j, s_0^j)\}_{j=1}^{n_p}$ from a logarithmic hypercube:
  $e_0^j \in I_e$, $s_0^j \in I_s$, $\forall j \in \{1, \ldots, n_p\}$
**for** $j \leftarrow 1$ **to** $n_p$ **do**
  $e_0 \leftarrow e_0^j$
  $s_0 \leftarrow s_0^j$
  $V_{\max} \leftarrow k_{\text{cat}} \cdot e_0$
  $C_s \leftarrow K_M \cdot \log(s_0) + s_0$
  Solve $\dot{s}^a(t) = -\frac{V_{\max}}{K_M + s^a(t)}$ on $[0, t_f]$
  Solve $c^a(t) = c_0 e^{-k_{\text{cat}}(t-t_0)} - s^a + s_0 e^{-k_{\text{cat}}(t-t_0)} + k_{\text{cat}} \int_0^{t_f} s^a(\tau) e^{-k_{\text{cat}}(t-\tau)} d\tau$ on $[0, t_f]$
  $t_{\max} \leftarrow_{t \in [0, t_f]} |s^a(t) - \frac{\gamma}{100} s_0|$
  $t_{\min} \leftarrow_{t \in [0, t_{\max}]} \dot{c^a}(t)$
  $P \leftarrow s^a - s_0$ on $[t_{\min}, t_{\text{fin}}]$
  $Q \leftarrow \int_{t_{\min}}^{t_{\text{fin}}} s^a(\tau) \cdot (e_0 - c^a(\tau))\, d\tau$
  $J \leftarrow -\int_{t_{\min}}^{t_{\max}} c^a(\tau)\, d\tau$
  Solve $(\bar{k_f}^j, \bar{k_r}^j) =_{(k_f, k_r) \in \mathbb{R}^2_{>0}} \big(P + Q \cdot k_f - R \cdot k_r\big)^2 + \lambda \big(K_m\, k_f - k_r - k_{\text{cat}}\big)^2$
**end**
$k_f^{\text{rec}} \leftarrow \prod_{j=1}^{n_p} \bar{k_f}^j$
$k_r^{\text{rec}} \leftarrow \prod_{j=1}^{n_p} \bar{k_r}^j$

---

### SI2. Local sensitivity analysis: sensitivity functions computation.

Consider any solution $s(t)$, $c(t)$ of the system (7)-(8). The corresponding local sensitivities, i.e. the partial derivatives of $s$ and $c$ with respect to $k_f$ and $k_r$, are evaluated as solutions of a system of ODEs. This system is obtained by the observation that, e. g.,

$$\frac{d}{dt}\frac{\partial s}{\partial k_f} = \frac{\partial}{\partial k_f}\frac{ds}{dt},$$

followed by replacement of $ds/dt$ with the right side of (7), and evaluation of the derivative. Similar remarks hold for the other sensitivities. A detailed development of the explicit calculations leads to the following linear system of 4 ODEs for 4 unknowns

$$\frac{d}{dt}\frac{\partial s}{\partial k_f} = -k_f\,(e_T - c)\,\frac{\partial s}{\partial k_f} + (k_f\,s + k_r)\,\frac{\partial c}{\partial k_f} - s\,(e_T - c) \tag{1}$$

$$\frac{d}{dt}\frac{\partial s}{\partial k_r} = -k_f\,(e_T - c)\,\frac{\partial s}{\partial k_r} + (k_f\,s + k_r)\,\frac{\partial c}{\partial k_r} + c \tag{2}$$

$$\frac{d}{dt}\frac{\partial c}{\partial k_f} = k_f\,(e_T - c)\,\frac{\partial s}{\partial k_f} - (k_f\,s + k_r + k_{\text{cat}})\,\frac{\partial c}{\partial k_f} + s\,(e_T - c) \tag{3}$$

$$\frac{d}{dt}\frac{\partial c}{\partial k_r} = k_f\,(e_T - c)\,\frac{\partial s}{\partial k_r} - (k_f\,s + k_r + k_{\text{cat}})\,\frac{\partial c}{\partial k_r} - c \tag{4}$$

with vanishing initial values.

### SI3. Minimum of the cost function $\mathcal{F}$.

Let's show now that the function $\mathcal{F}(k_f, k_r)$ introduced in (20) has a unique stationary point of coordinates $(\bar{k}_f, \bar{k}_r)$ given by (23).
The conditions for a minimum are given by $A_{ff} > 0$ and

$$\Delta = \frac{\partial^2 \mathcal{F}}{\partial k_f^2}\,\frac{\partial^2 \mathcal{F}}{\partial k_r^2} - \left(\frac{\partial^2 \mathcal{F}}{\partial k_f \partial k_r}\right)^2 = 4(A_{ff}\,A_{rr} - A_{fr}^2) > 0$$

at $(\bar{k}_f, \bar{k}_r)$.
With reference to the definitions of $A_{ff}$, $A_{rr}$, and $A_{fr}$, it is immediately seen that the inequality $A_{ff} > 0$ is always satisfied. Moreover, as to the sign of $\Delta$, it is found that

$$\Delta/4 = (\sum_i Q_i^2 + \lambda\,K_m^2)\,(\sum_i R_i^2 + \lambda) - (\sum_i Q_i R_i - \lambda\,K_m)^2,$$

whence it follows that

$$\Delta/4 = \sum_i Q_i^2 \sum_i R_i^2 - (\sum_i Q_i R_i)^2 + \lambda \sum_i (Q_i + K_m R_i)^2 \geqslant 0,$$

since $\sum_i Q_i^2 \sum_i R_i^2 - (\sum_i Q_i R_i)^2 \geq 0$ by the Cauchy-Schwarz inequality, $\lambda > 0$, and the sum of squares is necessarily non-negative.

Next, it can be observed that $\Delta/4 = 0$ if and only if for every $i \in \{1, ..., n\}$ either $Q_i = 0$ or $R_i = 0$ and at the same time $Q_i = -K_m R_i$; this means that both $\mathbf{Q}$ and $\mathbf{R}$ should be null.

### SI4. Single reconstructions.

Tables 1, 2, and 3 report the results of the 10 simulations performed for three representative reconstructions of $k_f$ and $k_r$, corresponding to cases no. 6, 10, and 12 in Table 2 of the manuscript. The structure of these tables follows the same format as Table 1.

| Simul. | $\mathbf{s_0}$ (nM) | $\mathbf{e_0}$ (nM) | $\mathbf{k_f^{rec}}$ (nM$^{-1}$ s$^{-1}$) | $\mathbf{k_r^{rec}}$ (s$^{-1}$) | $\frac{\mathbf{k_f^{rec}}}{\mathbf{k_f}}$ | $\frac{\mathbf{k_r^{rec}}}{\mathbf{k_r}}$ |
|---|---|---|---|---|---|---|
| 1 | 8.9284 | 1.7741 | 0.0457 | 0.4802 | 1.6345 | 64.2412 |
| 2 | 33.2603 | 1.0408 | 0.0425 | 0.4033 | 1.5179 | 53.9535 |
| 3 | 36.4052 | 2.7679 | 0.0411 | 0.3398 | 1.4678 | 45.4599 |
| 4 | 18.0640 | 6.1774 | 0.0390 | 0.2466 | 1.3939 | 32.9903 |
| 5 | 15.4046 | 7.9363 | 0.0352 | 0.1462 | 1.2592 | 19.5616 |
| 6 | 39.6392 | 3.8832 | 0.0404 | 0.3068 | 1.4432 | 41.0421 |
| 7 | 11.1903 | 4.4081 | 0.0397 | 0.2852 | 1.4194 | 38.1561 |
| 8 | 22.2301 | 7.4527 | 0.0382 | 0.2146 | 1.3659 | 28.7129 |
| 9 | 29.8209 | 5.7982 | 0.0403 | 0.2799 | 1.4391 | 37.4447 |
| 10 | 24.4240 | 8.9637 | 0.0368 | 0.1701 | 1.3166 | 22.7521 |
| G. mean | - | - | 0.0398 | 0.2708 | 1.4223 | 36.2288 |

Table 1: Reconstruction no. 6. $K_m = 28.47\,\text{nM}$, $k_{\text{cat}} = 0.7893\,\text{s}^{-1}$.

| Simul. | $\mathbf{s_0}$ (nM) | $\mathbf{e_0}$ (nM) | $\mathbf{k_f^{rec}}$ (nM$^{-1}$ s$^{-1}$) | $\mathbf{k_r^{rec}}$ (s$^{-1}$) | $\frac{\mathbf{k_f^{rec}}}{\mathbf{k_f}}$ | $\frac{\mathbf{k_r^{rec}}}{\mathbf{k_r}}$ |
|---|---|---|---|---|---|---|
| 1 | 5640.8365 | 2076.4856 | 0.0008 | 1.1823 | 1.3696 | 12.7167 |
| 2 | 6769.7493 | 2449.3611 | 0.0008 | 1.0419 | 1.3493 | 11.2065 |
| 3 | 1801.6283 | 1744.8800 | 0.0007 | 0.7596 | 1.2172 | 8.1707 |
| 4 | 10679.1356 | 873.9609 | 0.0008 | 1.8910 | 1.4868 | 20.3404 |
| 5 | 5189.7409 | 575.4829 | 0.0009 | 2.5133 | 1.6214 | 27.0339 |
| 6 | 3352.0249 | 189.0155 | 0.0031 | 18.9851 | 5.5823 | 204.2097 |
| 7 | 3500.1283 | 2013.8304 | 0.0007 | 0.9521 | 1.2924 | 10.2407 |
| 8 | 7545.0421 | 1272.6103 | 0.0008 | 1.7068 | 1.4669 | 18.3589 |
| 9 | 9548.2622 | 927.5489 | 0.0008 | 1.8740 | 1.4862 | 20.1576 |
| 10 | 9162.3688 | 1384.0429 | 0.0008 | 1.6093 | 1.4484 | 17.3103 |
| G. mean | - | - | 0.0009 | 1.8267 | 1.6187 | 19.6488 |

Table 2: Reconstruction no. 10. $K_m = 7541.26\,\text{nM}$, $k_{\text{cat}} = 4.1564\,\text{s}^{-1}$.

| Simul. | $\mathbf{s_0}$ (nM) | $\mathbf{e_0}$ (nM) | $\mathbf{k_f^{rec}}$ ($\mathrm{nM}^{-1}\ \mathrm{s}^{-1}$) | $\mathbf{k_r^{rec}}$ ($\mathrm{s}^{-1}$) | $\frac{\mathbf{k_f^{rec}}}{\mathbf{k_f}}$ | $\frac{\mathbf{k_r^{rec}}}{\mathbf{k_r}}$ |
|---|---|---|---|---|---|---|
| 1 | 48.5273 | 14.2518 | 0.0808 | 0.0010 | 0.8986 | 0.0015 |
| 2 | 76.8753 | 16.8140 | 0.0827 | 0.0308 | 0.9204 | 0.0462 |
| 3 | 40.0074 | 12.8567 | 0.0804 | 0.0110 | 0.8948 | 0.0166 |
| 4 | 53.6288 | 2.8521 | 0.1226 | 2.2427 | 1.3642 | 3.3704 |
| 5 | 37.2667 | 11.6484 | 0.0799 | 0.0003 | 0.8884 | 0.0004 |
| 6 | 22.1992 | 9.9379 | 0.0798 | 0.0394 | 0.8877 | 0.0592 |
| 7 | 67.2890 | 4.8010 | 0.1017 | 1.1332 | 1.1317 | 1.7030 |
| 8 | 27.1128 | 7.7954 | 0.0848 | 0.2785 | 0.9437 | 0.4185 |
| 9 | 12.1706 | 3.0178 | 0.0813 | 0.1798 | 0.9043 | 0.2702 |
| 10 | 65.0324 | 6.8351 | 0.0964 | 0.8205 | 1.0724 | 1.2330 |
| G. mean | - | - | 0.0882 | 0.0577 | 0.9809 | 0.0868 |

Table 3: Reconstruction no. 12. $K_m = 52.73\,\mathrm{nM}$, $k_{\mathrm{cat}} = 4.0750\,\mathrm{s}^{-1}$.
